# Combining IP_3_ affinity chromatography and bioinformatics reveals a novel protein-IP_3_ binding site on *Plasmodium falciparum* MDR1 transporter

**DOI:** 10.1101/2021.03.25.437059

**Authors:** Eduardo Alves, Helder Nakaya, Euzébio Guimarães, Célia R. S. Garcia

## Abstract

Intracellular Ca^2+^ mobilization induced by second messenger IP_3_ controls many cellular events in most of the eukaryotic groups. Despite the increasing evidence of IP_3_-induced Ca^2+^ in apicomplexan parasites like *Plasmodium*, responsible for malaria infection, no protein with potential function as an IP_3_-receptor has been identified. The use of bioinformatic analyses based on previously known sequences of IP_3_-receptor failed to identify potential IP_3_-receptor candidates in any *Apicomplexa*. In this work, we combine the biochemical approach of an IP_3_ affinity chromatography column with bioinformatic meta-analyses to identified potential vital membrane proteins that present binding with IP_3_ in *Plasmodium falciparum*. Our analyses reveal that PF3D7_0523000, a gene that codes a transport protein associated with multidrug resistance, as a potential target for IP_3_. This work provides a new insight for probing potential candidates for IP_3_-receptor in *Apicomplexa*.

## Introduction

The inositol 1,4,5-triphosphate (IP_3_) is an important second messenger that regulates cytosolic Ca^2+^ in a variety of Eukaryotic organism^(1, 2)^. Briefly, the activation of phospholipase C (PLC) mediated by surface receptor breaks phosphatidylinositol 4,5-bisphosphate (PIP_2_) into soluble short life second messenger IP_3_ that binds into IP_3_ receptor (IP_3_R) culminating in intracellular Ca^2+^ release^(3, 4)^.

The phylum *Apicomplexa* includes unicellular eukaryotes parasites like *Plasmodium*, the etiology agent of malaria infection, and possesses the metabolic enzymes responsible for generation and degradation of IP_3_, see review ^(5)^. IP_3_ can mobilize Ca^2+^ from intracellular stores in isolate and permeabilize blood stage *P. chabaudi*^(66)^ and in intact *P. falciparum* within red blood cells (RBCs). Within RBCs, parasites manage to maintain the Ca stores full even under low Ca^2+^ environment^(8)^. An increasing number of reports supporting the existence of intracellular Ca^2+^ release induced by IP_3_ in malaria parasites^(6, 7, 9–12)^ suggesting the existence of a Ca^2+^ channel sensitive to IP_3_, the IP_3_R.

The IP_3_R it is a well know described protein in vertebrates that contains around four to six transmembrane domains (TMDs), see review^(13)^. Prole and Taylor^(14)^ used the sequence of mammal N-terminal IP_3_R binding domain and the amino-terminal RIH (Ryanodine and IP_3_R homology) domains to perform a BLAST (Basic Local Alignment Search Tool) on the genome of diverse parasites. However, this work failed to find any potential candidate for IP_3_R in *Apicomplexa*. So far, no apicomplexan IP_3_R candidate has been identified through bioinformatics approaches. Moreover, there is no publication that attempted to use biochemical approach like an IP_3_ affinity chromatography column in *Apicomplexa* to identify proteins that might bind to IP_3_.

Hirata and collaborators^(15)^ managed to enrich proteins from rat brain sample that has an affinity to IP_3_ like IP_5_-phosphatase and IP_3_ 3-kinase using an analogous IP_3_ affinity chromatography column 2-O-[4-(5-aminoethul-2-hydroxyphenylazo) benzoyl]−1,4,5-tri-O-phosphono-myo-inositol trisodium salt-Sepharose 4B. Using a similar column, Kishigami and collaborators^(16)^ managed to identify the PLC protein from octopus’ eyes *Todarodes pacificus* and reported that squid rhodopsin also has an affinity to IP_3_. Nevertheless, besides the potential of these columns to enrich proteins that bind to IP_3_, no IP_3_R has ever been identified using a IP_3_-affinity column alone.

By adapting the protocol from Hirata/Kishigami^(15, 16)^, we created a column containing IP_3_ conjugated with biotin linked with a high-performance sepharose-streptavidin and challenged with proteins from asynchronous *P. falciparum* blood-stage. Using a IP_3_-free column containing only sepharose-streptavidin as reference, we selected only the candidates exclusive on the IP_3_-column to undergo a series of bioinformatic meta-analyses. Our approach targeted for candidates with at least one transmembrane domain, considered essential, conserved among most apicomplexan species, and with unknown or nor-clear function. Finally, the candidate that fit all these criteria were used as targets for *in silico* molecular docking against IP_3_.

Using this strategy, we identified the *P. falciparum* multidrug resistance protein 1 (*Pf*MDR1), a vital and conserved membrane protein that has potential to bind to IP_3_. This protein is located on parasite food vacuole, a Ca^2+^ storage compartment^(17)^. Combined, the affinity column and bioinformatic successfully provided a candidate for a membrane protein associated to an intracellular Ca^2+^ compartment that binds to IP_3_ in malaria.

## Material and Methods

### *P. falciparum* Culture

*P. falciparum* (D37) parasites were maintained in culture as described^(18)^. Briefly, *P. falciparum* were cultured in RPMI media supplemented with 50 mg/L hypoxanthine; 40 mg/L gentamycin; 435 mg/L NaHCO_3_; 2% haematocrit of A^+^ human red blood cells and 10% A^+^ human blood serum in an atmosphere of 5% CO_2_; 3% O_2_; 92% N_2_ at 37°C. Media was changed every 24 h and RBCs replaced every 48 h. Parasitemia and the development stage of cultures were determined by Giemsa-stained smears.

### *P. falciparum* protein sample

Total *P. falciparum* protein extract was obtained from 2.5 L of unsynchronized culture, at 8% parasitemia. Culture was washed three times in PBS (300 g, 5 minutes) and parasites isolated from erythrocytes using 0.03% (w/v) saponin (Sigma) on PBS containing protease inhibitors: antiplaque, pepstatin, chymostatin and leupeptin (Sigma) at concentrations of 20 μg/mL each and 500 μM benzamidine (Sigma). Isolate parasites were centrifuge on 1300 g for 10 minutes at 4°C and washed three times in PBS with protease inhibitors. The isolate parasite samples were resuspended in 50 mM TRIS-HCl buffer pH 7.4 containing 2 mM EDTA, 0.1% Triton X−100, protease inhibitors and 1 mM PMSF. Samples were sonicated on SONIC (Vibracell) 50% potency for 20 seconds for 3 times on ice (10-second interval between each sonication) follow by at 1300 g centrifugation for 10 minutes at 4°C for removal of the insoluble pellet. DNAse and RNAse (final concentration 200 ng/μL each) were added on soluble pellet and incubate for one hour at 37 °C. The samples were passed through a 0.45 μm filter. The amount of protein was quantitated using Pierce’s BCA protein assay kit.

### IP_3_-affïnity chromatography column

To build the column, it was used a commercial high performance Sepharose substrate bound to streptavidin (GE Healthacare Life Science) and biotin-conjugated IP_3_ (Echelom Biosciences). The streptavidin-sepharose column was equilibrated by washing once with 10x volume of ice-cold, 0.45 μm filter binding buffer (20 mM NaH_2_PO_4_, 150 mM NaCl, 20 mM LiCl and 2 mM EDTA, pH 7.5). The columns were mounted in a 15 mL sterile falcon tube. For each column it was used 1.25 ml of equilibrated S epharos e-streptavidin resuspended in binding buffer mixed with 20 μg of IP_3_-biotin. The columns were left by constant stirring for 12 hours at 4°C in a dark environment and then centrifuged for 1 minute, 300 g at 4 °C. The supernatant containing excess IP_3_-biotin was removed and columns were washed five times with 2 ml of ice-cold binding buffer to remove any free IP_3_-biotin. Two distinct columns were assembled: one containing IP_3_-biotin-sepharose-streptavidin and other containing sepharose-streptavidin only. In each column was added 2.5 mg of *P. falciparum* protein extract. The column volume was adjusted with ice-cold binding buffer with protease inhibitors until a final volume of 5 ml. The columns were incubated at 4°C under gentle, steady shaking in a light-protected environment for 12 hours. After incubation, the columns were centrifuged for 1 minute at 300 g at 4 °C and the supernatant discarded. Each column was washed seven times with ice-bound binding buffer with protease inhibitors. To elute the proteins, 1mL of an ice-cold elution buffer (8 M Guanidin-HCl, 20 mM LiCl, 2 mM EDTA, pH: 1.5 with protease inhibitors) was added on each column and followed by constant stirring for 1 hour at 4 °C. At the end of this incubation the columns were centrifuged for 1 minute at 300 g and the supernatant collected in sterile low binding protein eppendorfs.

### Mass spectrometry

The protein samples were applied on 8% polyacrylamide gel and run at low voltage (60 v) until the bands were discriminated. After the run, the gel was fixed and stained following the recommendations of the “Colloidal Blue Staining Kit” from Invitrogen. The sections of the gel containing visible bands were cut and sent for analysis on a mass spectrometer at Taplin Mass Spectrometry, Harvard Medical School (https://taplin.med.harvard.edu/) for protein identification. All identified proteins containing at least one exclusive peptide match was considered for analyses.

### Transmembrane domain prediction

To detect the presence of a transmembrane domain, the whole amino acid sequence from the protein identified at mass spectrometry were analysed using the public HMMTOP program version 2.0 (www.enzim.hu/hmmtop/). This program predicts the number of transmembrane helicases and their position from peptide/protein amino acid sequence.

### Phenotype score, conservation and function predictions

The phenotype score used the determinate gene essentiality of each candidate was obtained from the work of Zhang and collaborators^(19)^ that is available on PlasmoDB (https://plasmodb.org/plasmo). To identify the presence of a candidate orthologs among *Apicomplexa* group, we use the OrthoMCL database (https://orthomcl.org/orthomcl). For function prediction, we consulted the gene annotation information provided by PlasmoDB.

### *In silico* docking with IP_3_

The primary sequence of MDR1 (Gene - PF3D7_0523000, plasmodb.org) was used to build its probable 3D structure by homology modelling. The server SwissModel^(20)^ was employed to automatically build the models optimized to bind IP_3_ at various locations inside MDR1 homology. Blind molecular docking simulation were carried out to obtain possible interactions for the intermembrane domain as predicted by the TMHMM Server^(21)^. The SwissDock^(22)^ server enabled the study of IP_3_ intermembrane MDR1 domain binding poses. Additionally, the IP_3_-Ion-MDR1 binding was further investigated using the multidrug transporter permeability (P)-glycoprotein is an adenosine triphosphate (ATP)-binding cassette (PDB id: 6C0V). The later ability to bind simultaneously ATP and a divalent cation at the intracellular domain was used to guide the inspect a hypothetical IP_3_-Ion-MDR1 interaction. IP_3_ was manually positioned inside the ATP cavity to mimic an IP_3_-Mg^2+^ interaction. The binding conformation was optimized with molecular mechanics by means of the UCSF^(23)^ chimera minimize structure tools.

### Protein-protein interaction network

Using *Plasmodium* interactome data^(24)^, we looked for the proteins that interact with the MDR1. The protein annotation and functions were also retrieved from the original publication. Network was generated using Cytoscape^(25)^.

## Results

### IP_3_-affinity chromatography data

Adapting the protocol based on Hirata/Mishigami^(15, 16)^, we use an IP_3_ affinity chromatography column with protein homogenate from unsynchronized asexual blood stages of isolated *P. falciparum* as the first step to identified potential proteins that have a similar function to IP_3_R receptor in a mammal.

The access code of the brute data on mass spectrometry analyses from the eluate samples of IP_3_-affinity chromatography column can be found in Supplemental Material Table 1. At least 700 proteins from *P. falciparum* containing at least one exclusive peptide were detected from the IP_3_-sepharose column. In comparison, 494 proteins were detected from the sepharose matrix alone (Figure 1). With both columns’ information, all proteins exclusively present on IP_3_-sepharose were selected (total 206 proteins) for the bioinformatic meta-analyses (Sup. Table 2).

**Figure 1:**
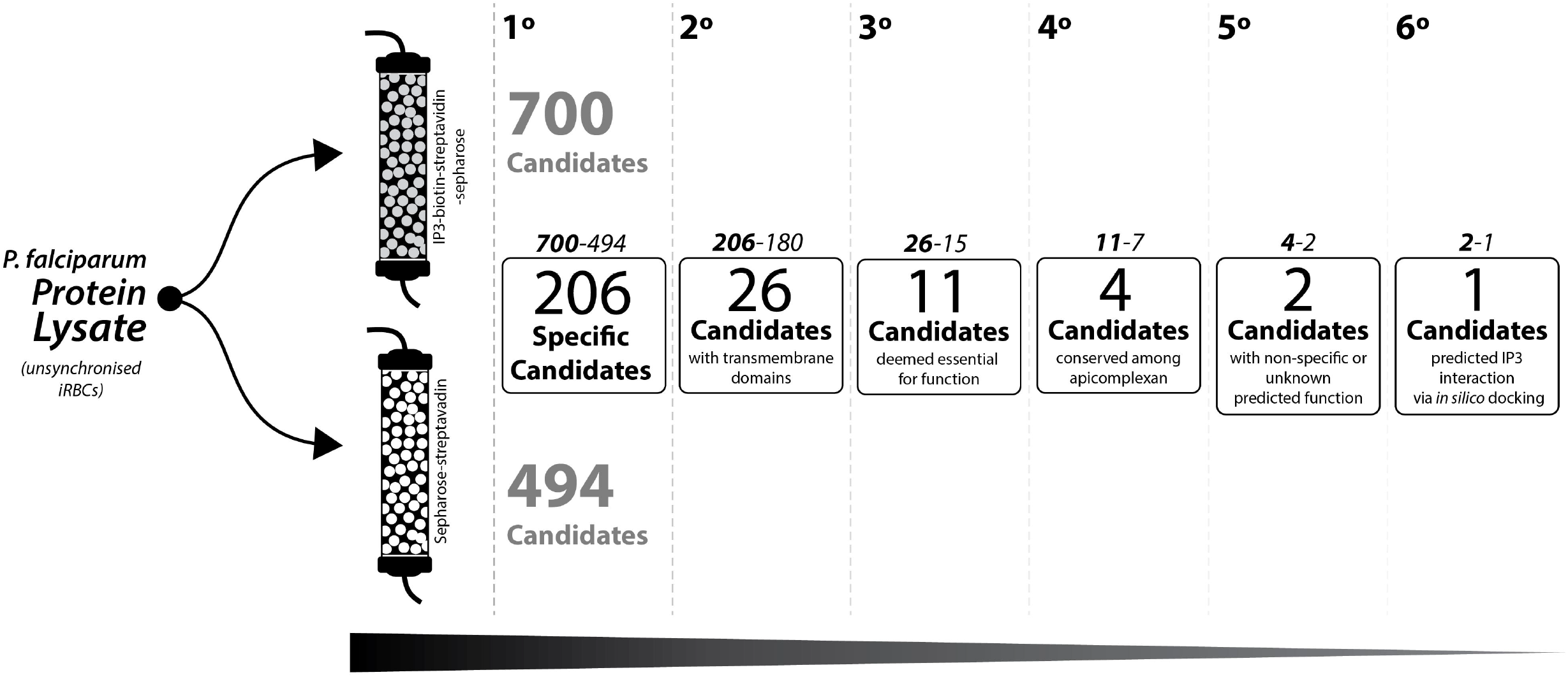
Schematic approached to pinpoint potential candidates for IP_3_R from isolates blood stage *P. falciparum*. 1° step: selection of proteins that are exclusively found on IP_3_-Biotin-streptavidin-sepharose column. 2° step: selection of proteins that contains at least one transmembrane domain. 3° step: selection of protein that are considered essential for malaria parasite during red blood stage development. 4° step: Selection of proteins that are conserve among most species within *Apicomplexa* group. 5° step: selection of protein with unknow or non-specific metabolic function. 6° step: candidates with positive *in silico* docking against IP_3_.

**The table I:**
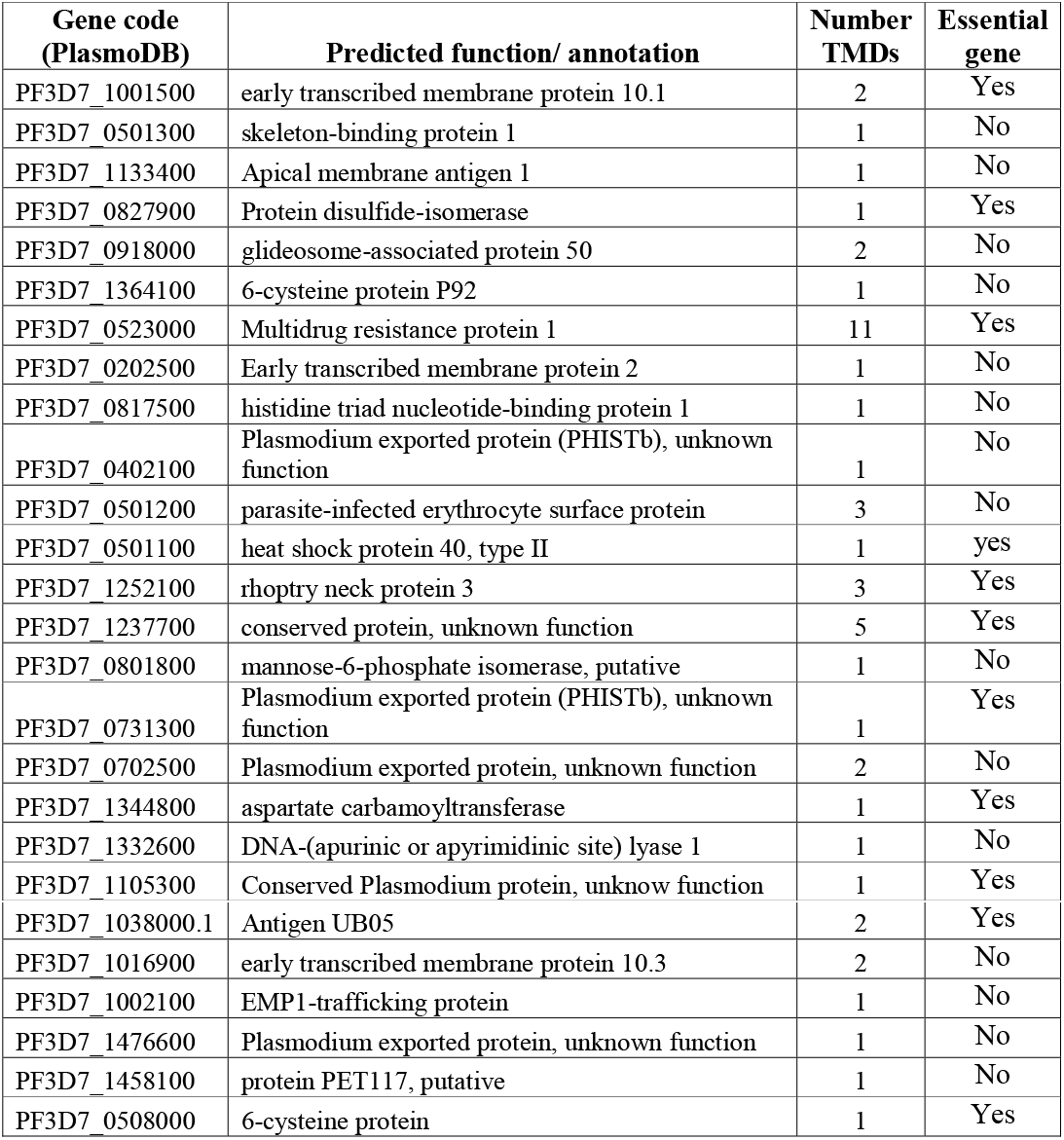
The list of 26 proteins exclusively found at IP_3_-biotin-streptavidin-sepharose column that contains at least one TMDs.

| Gene code<br>(PlasmoDB) | Predicted function/ annotation | Number<br>TMDs | Essential<br>gene |
| --- | --- | --- | --- |
| PF3D7_1001500 | early transcribed membrane protein 10.1 | 2 | Yes |
| PF3D7_0501300 | skeleton-binding protein 1 | 1 | No |
| PF3D7_1133400 | Apical membrane antigen 1 | 1 | No |
| PF3D7_0827900 | Protein disulfide-isomerase | 1 | Yes |
| PF3D7_0918000 | glideosome-associated protein 50 | 2 | No |
| PF3D7_1364100 | 6-cysteine protein P92 | 1 | No |
| PF3D7_0523000 | Multidrug resistance protein 1 | 11 | Yes |
| PF3D7_0202500 | Early transcribed membrane protein 2 | 1 | No |
| PF3D7_0817500 | histidine triad nucleotide-binding protein 1 | 1 | No |
| PF3D7_0402100 | Plasmodium exported protein (PHISTb), unknown function | 1 | No |
| PF3D7_0501200 | parasite-infected erythrocyte surface protein | 3 | No |
| PF3D7_0501100 | heat shock protein 40, type II | 1 | yes |
| PF3D7_1252100 | rhoptry neck protein 3 | 3 | Yes |
| PF3D7_1237700 | conserved protein, unknown function | 5 | Yes |
| PF3D7_0801800 | mannose-6-phosphate isomerase, putative | 1 | No |
| PF3D7_0731300 | Plasmodium exported protein (PHISTb), unknown function | 1 | Yes |
| PF3D7_0702500 | Plasmodium exported protein, unknown function | 2 | No |
| PF3D7_1344800 | aspartate carbamoyltransferase | 1 | Yes |
| PF3D7_1332600 | DNA-(apurinic or apyrimidinic site) lyase 1 | 1 | No |
| PF3D7_1105300 | Conserved Plasmodium protein, unknow function | 1 | Yes |
| PF3D7_1038000.1 | Antigen UB05 | 2 | Yes |
| PF3D7_1016900 | early transcribed membrane protein 10.3 | 2 | No |
| PF3D7_1002100 | EMP1-trafficking protein | 1 | No |
| PF3D7_1476600 | Plasmodium exported protein, unknown function | 1 | No |
| PF3D7_1458100 | protein PET117, putative | 1 | No |
| PF3D7_0508000 | 6-cysteine protein | 1 | Yes |

Once the proteins exclusive for IP_3_-column were identified (Sup. Table 2), the first bioinformatic approach aimed to select proteins that contains a transmembrane domain (TMD). The IP_3_R is a protein attached at the membrane, a TMD is an important structure to anchor proteins through biological membranes by its physical properties like the length and hydrophilicity of the transmembrane span^(26)^, every IP_3_R in vertebrates, invertebrates and single eukaryotes organism possess a TMDs, so we use this feature as the second step to select potential candidates for IP_3_R. Figure 1.

The table 1 summarizes the list of 26 proteins exclusively found at IP_3_-biotin-streptavidin-sepharose column that contains at least one TMDs. Transfection of *P. falciparum* to constitutively express IP_3_-sponge, a protein containing a modified IP_3_ binding domain based on mouse IP_3_R that sequestrate cytosolic IP_3_^(27)^, does not result in viable parasites^(28)^ suggesting a vital role of IP_3_ signalling in *P. falciparum*. With this information, the next step to narrow the number of potential candidates that might act as IP_3_R in malaria is focus on essential genes. To deem whether a gene is essential, we considered only the genes that score lower than 0.5 on its mutagenic index of phenotype graphic (data provide by PlasmoDB). That decrease the number of candidates to 11 (Fig. 1, table I).

For the next step on the bioinformatic meta-analyses, we considered only the genes that is conserve among multiples species within *Apicomplexa* phylum. Only four essential candidates with TMDs domains met this criterium: multidrug resistance protein 1 (MDR1); a heat shock protein 40, type II (HSP40); aspartate carbamoyltransferase (ATCase) and antigen UB05. PlasmoDB access code: PF3D7_0523000, PF3D7_0501100, PF3D7_1344800 and PF3D7_1038000 respectively. Among these 4 candidates, only MDR1 and antigen UB05 has unknow or unclear function. The HSP40 is a cochaperone protein with conserved J-domain that regulates other heat socks protein 70 (HSP70)^(29)^ and the ATCase is an enzyme important for the pyrimidine biosynthetic pathway^(30)^.

### IP3-MRD1 binding modelling and protein interactions network

The MDR1 model provide by the SwissModel server proved to be quite similar to human P-glycoprotein ABCB1 receptor, protein data bank id: 7A69^(31)^. The sequence alignment proved that a homology model could be built with fair quality with an identity of 29.7% and similarity of 48.2% (Pairwise Sequence Alignment EMBOSS Water server, https://www.ebi.ac.uk/Tools/emboss/). Two binding position at the transmembrane domain of MRD1 and IP_3_ binding was estimated by the SwissDock server (Figure 2).

**Figure 2:**
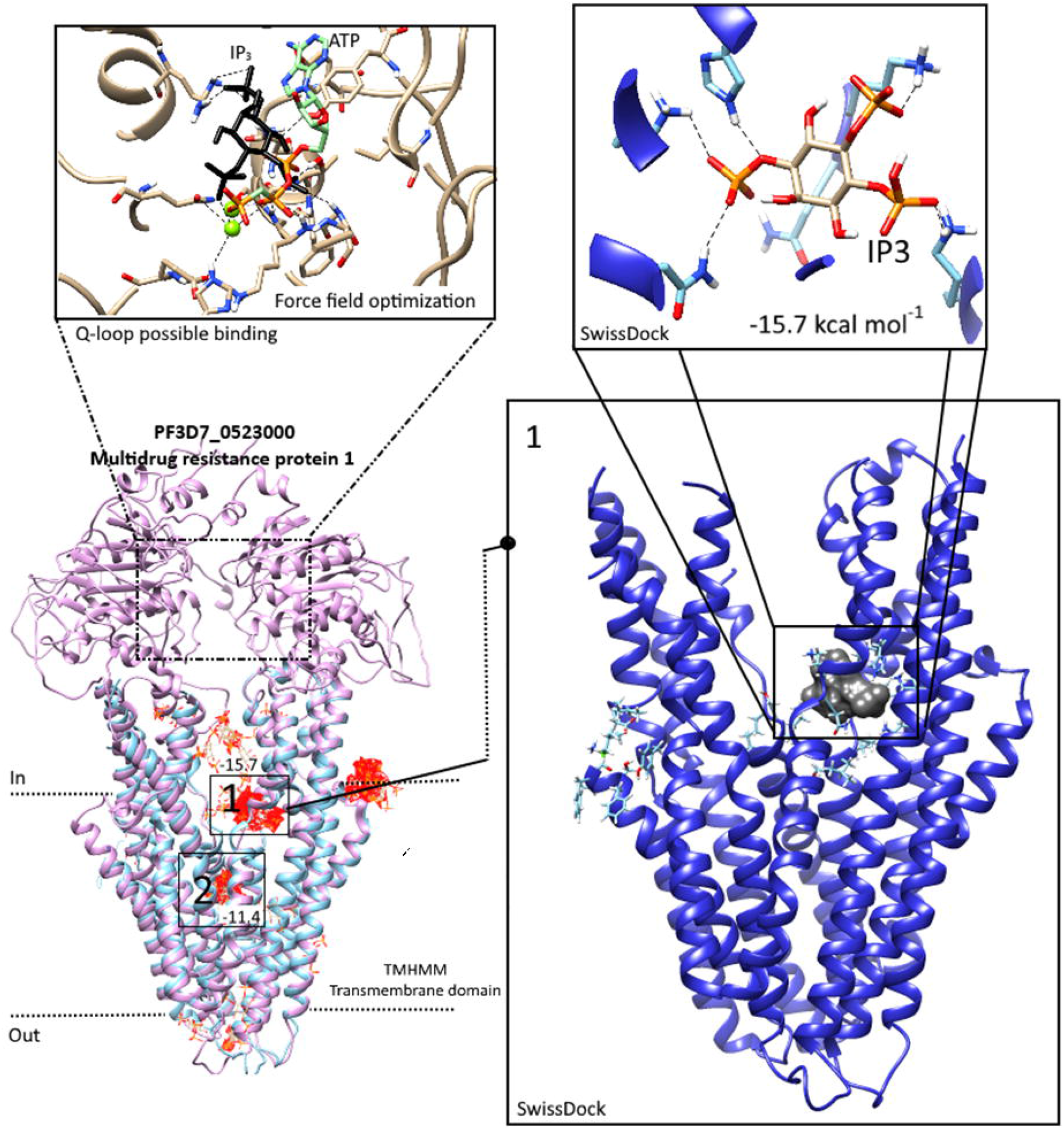
Schematic representation of the SwissDock most energetically favored binding poses of IP_3_-MDR1. The cytosolic nucleotide-binding domain (upper part) display an IP_3_ associated with divalent cation (green spheres) interacting on the same ATP binding pocket. The transmembrane domains (lower part) display two possible pocket sites on IP_3_ interaction and their respective interaction energy values (kcal/mol). On the right side, details of the IP_3_-MDR1 pocket 1 interaction, a region rich on lysine.

The pocket 1 (binding energy −15.7 kcal/mol) proved to be the best IP_3_ docking position. The site is a lysin rich domain able to form various hydrogen bonds with IP_3_. The second-best bind pocket proved to be less favored as derived from the lower interaction energy (−11.4 kcal/mol). Another binding possibility investigated was the interaction with the same pocket ATP binding. The interaction involves the presence of a divalent cation (green spheres) like Mg^2+^ intercalating with IP_3_. The MDR1 is an ATP-binding cassette (ABC) transporter family member that is associated to multidrug drug resistance due the ability to translocating amphiphilic compounds^(32)^. The translocation of a substrate across the membrane by proteins like *P. falciparum* MDR1 requires an ATP binding on Q-loop site that cause a rearrangement of TM^(33)^. The binding on IP_3_-divalent cation on the MDR1 Q-loop site suggests a potential competition between ATP and IP_3_. Interestingly, ATP is known to allosterically modulate the functional of mammal IP_3_R^(34, 35)^ including the inhibition of Ca^2+^ flux regulated by IP_3_R under high concentration of ATP^(35)^.

To help uncover the cellular function of MDR1 protein, we searched for proteins that interact with MDR1 in the *Plasmodium* interactome data^(24)^ (Figure 3). It is interesting to note that MDR1 interacts with receptor for activated C kinase (RACK1, PF3D7_1148000). The *Pf*RACK1 can inhibit host IP_3_-mediated Ca^2+^ signaling by direct interaction with IP_3_R(36). This data suggests that *Pf*MDR1 has the potential to associate or have similar functions to other receptors that coordinate signaling events regulated by protein kinase C.

**Figure 3:**
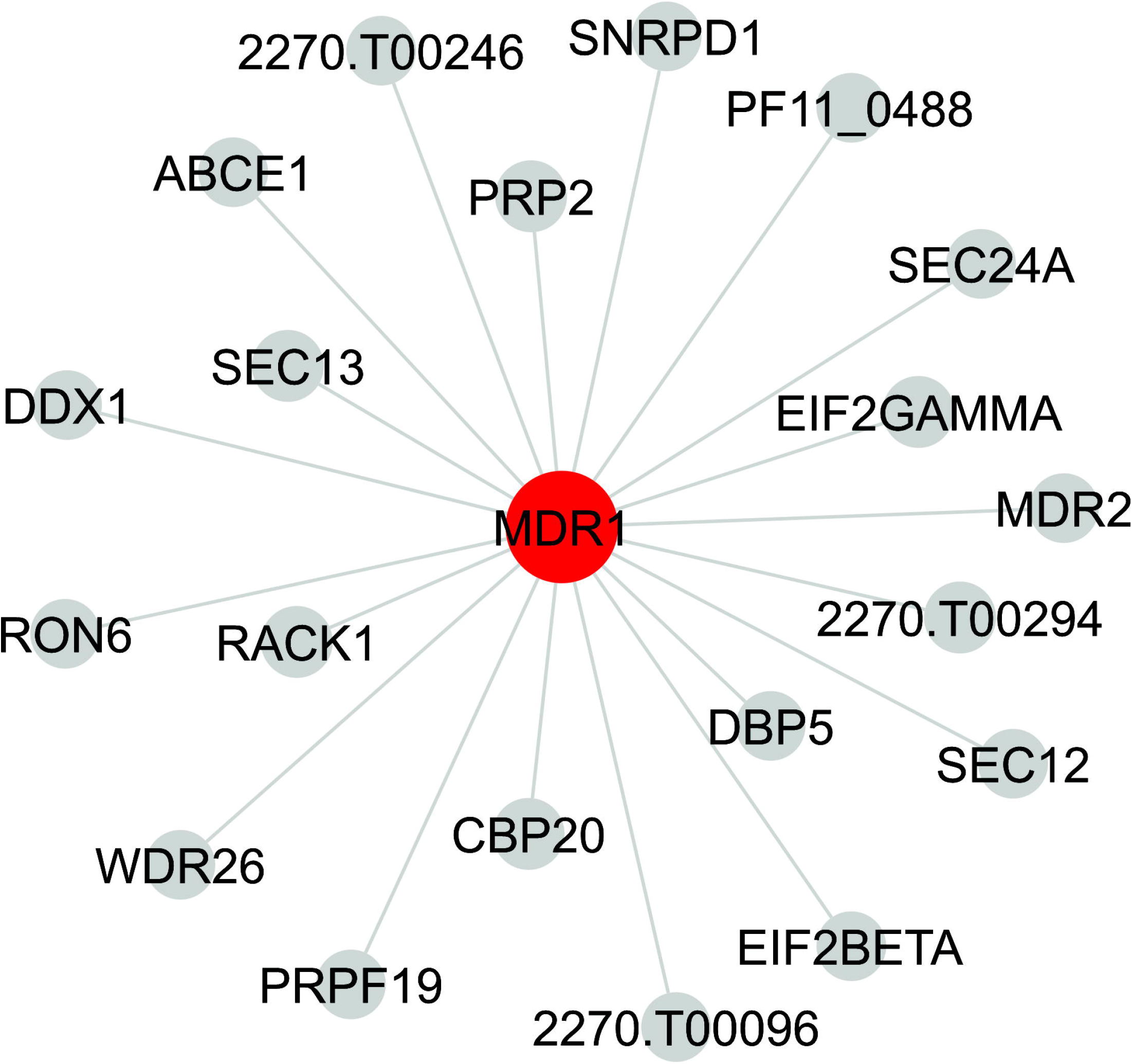
Proteins that interact with MDR1. The network shows the proteins (grey nodes) that interact with MDR1 (red node) according to the *Plasmodium* interactome data^(24)^. Gene codes: 2270.T00246, translation initiation factor (PF3D7_0607000). SNRPD1, small nuclear ribonucleoprotein Sm D1(PF3D7_1125500). ABCE1, ABC transporter E family member 1 (PF3D7_1368200). PRP2, pre-mRNA-splicing factor ATP-dependent RNA helicase PRP2 (PF3D7_1231600). PF11_0488, serine/threonine protein kinase (PF3D7_1148000). DDX1, ATP-dependent RNA helicase DDX1 (PF3D7_0521700). SEC13, protein transport protein SEC13 (PF3D7_1230700). SEC24A, transport protein Sec24A (PF3D7_1361100). EIF2GAMMA, eukaryotic translation initiation factor 2 subunit gamma (PF3D7_1410600). RON6, rhoptry neck protein 6 (PF3D7_0214900). RACK1, receptor for activated C kinase (PF3D7_0826700). MDR2, multidrug resistance protein 2 (PF3D7_11447900). 2270.T00294, ATP-dependent RNA helicase MTR4, (PF3D7_0602100). DBP5, ATP-dependent RNA helicase DBP5 (PF3D7_1459000). SEC12, guanine nucleotide-exchange factor (PF3D7_1116400). WDR26, WD repeat-containing protein 26 (PF3D7_0518600). CBP20, nuclear cap-binding protein subunit 2 (PF3D7_0415500). PRPF19, pre-mRNA-processing factor 19 (PF3D7_0308600). EIF2BETA, eukaryotic translation initiation factor 2 subunit beta (PF3D7_1010600). 2270.T00096, cleavage stimulation factor subunit 1 (PF3D7_0620500).

## Discussion and conclusion

Phylogenetic analyses and comparative genomic managed to reveal both unique and conserved proteins related to calcium signalling pathways on apicomplexan parasites^(14, 37, 38)^, nevertheless the IP_3_R still remain a major missing piece of this Ca^2+^ signalling toolkit. The conflict with the pharmacological and functional studies using exogenous IP_3_ that supports the existence of protein sensitive to IP_3_ that is capable to mobilize Ca^2+^ with the bioinformatic data that constantly failed to point an IP_3_R candidate suggests this receptor in *Apicomplexa* has a different and unique structure compared to the IP_3_-binding core domain from other eukaryotes. The search for an IP_3_R in *Apicomplexa* requires a different strategy that do not rely exclusively on bioinformatic tools as BLAST (Basic Local Alignment Search Tool) based on previously known IP_3_R.

The use of IP_3_ affinity chromatography column has being successfully reported to concentrate proteins that interact with high affinity to IP_3_ analogues^(15)^ as well as retained key components from IP_3_-Ca^2+^ signalling from proteins extract from tissues^(16)^. In this work, we used a biotin-inositol 1,4,5-triphosphate attached to a high-performance streptavidin-sepharose substrate to initially enrich proteins with IP_3_ affinity from unsynchronized isolate *P. falciparum* blood culture. One of the major limitations of using chromatography affinity column based on a short life IP_3_ molecule is the number of proteins that are naturally present at sample homogenate that degrades this second messenger. *P. falciparum* contains proteins that can dephosphorylate or phosphorylate IP_3_ like inositolpolyphosphate 5-phosphatase and inositol 1,4,5-trisphosphate 3-kinase^(39)^. In this protocol, we tried to overcome this limitation by keeping all the binding and elution steps under low temperature while adding LiCl in every buffer. LiCl has being previously used to inhibit the dephosphorylation of IP_3_ ^(40–42)^. Another risk of using an IP_3_ affinity column is assume that protein(s) that might act as IP_3_R in *Plasmodium* do not bind/interact with strong affinity with the sepharose-streptavidin substrate alone. In this protocol, we excluded all 494 proteins that bind with the sepharose-free IP_3_ column as a potential IP_3_R candidate (Fig.1).

From the 206 proteins identified exclusively from IP_3_-sepharose column, 180 did not contained any TMDs suggesting the protocol used to extract the proteins from parasite lysate benefit mostly soluble proteins that do not strongly interact with lipid bilayers. For future trials, this protocol can be strongly optimized by using a protein extraction that target membrane proteins (MPs). The IP_3_R in mammals is a MP protein containing 6 TMDs^(43)^. The presence of a TMDs is an important aspect of any MPs to physically interact with biological membranes^(26)^. It is fair to predict that any protein with potential function of IP_3_R should have TMDs to able to interact with membranes. The table I list all candidates with TMDs exclusively from IP_3_-column.

Ca^2+^ is a second messenger that regulate a variety of vital functions in apicomplexan parasites^(37, 44, 45)^. Accordingly, the use of 2-aminoethoxydiphenyl borinate (2-APB), a pharmacological drug that inhibit IP_3_R, abolished spontaneous Ca^2+^ mobilization and compromise intracellular development of blood stage *P. falciparum*^(11^. Pecenin and collaborator^(28)^ failed to obtain any viable parasite expressing IP_3_-sponge. These data suggest that IP_3_-Ca^2+^ signalling pathway has a vital role during intraerythrocytic development of *P. falciparum* and support our hypotheses that a potential candidate for IP_3_R in *Plasmodium* not only has to present a TMDs, but also has to be essential. A prediction of gene essentiality in *P. falciparum*, based on the work of Zhang and collaborators^(19)^ is available for consultation at PlasmoDB website.

The pharmacological evidence that supports IP_3_-Ca^2+^ signalling pathway in *Apicomplexa* group is not exclusive to malaria parasites, but also present in *Toxoplasma gondii^46^’^48^* and *Babesia bovis*^(49)^. The strategy to pinpoint the potential candidate for IP_3_R in apicomplexan should not rely on gene only excusive to *Plasmodium* species. Adding this extra meta-analysis step, the list of potential candidates presented exclusively on IP_3_-sepharose column is finally reduced to four proteins: a MDR1; a HSP40); an ATCase and antigen UB05. Among those four, only MDR1 and antigen UB05 has an unclear function.

The small number of candidates makes the use of more computational demanding bioinformatic analyses more feasible. A molecular docking allows to target the structural protein complexes from our candidate list against potential ligand as IP_3_ or other potent IP_3_-analogues drugs like adenophostin A^(50)^.

Molecular docking on IP_3_ on *P. falciparum* MDR1 protein revealed two potential binding sites on TMD: pocket site **1** (binding energy −15.7 kcal/mol) and pocket site **2** (−11.4 kcal/mol), see Figure 2. This data suggests that MDR1 pocket **1** has a higher affinity to IP_3_ compared to IP_3_-binding core of mammal IP_3_R (ΔG= −10.3 kcal/mol on 23°C)^(51)^ and a lower affinity when compare to IP_3_-binding with N-terminal region of mammal IP_3_R (ΔG= −79.5 kcal/mol)^(52)^. Nevertheless, the binding of ATP on Q-Loop site on nucleotide-binding domain (NBD) likely cause profound changes on the TMD region^(33)^making hard to predict the real affinity of MDR1 protein with IP_3_.

In *P. falciparum* the MDR1 gene encodes for a 162.2 kilo Daltons P-glycoprotein located on the digestive vacuole (DV)^(53)^ with unclear function, but the polymorphisms within this protein are associated to increase *in vitro* resistance against multiple antimalarial drugs like quinine^(54–61)^. The MDR1 display a role as a transporter protein bringing solutes into DV and it is consisted of two distinct homologous regions: one cytosolic nucleotide-binding domain (NBD) and a substrate-binding consisting of 11 TMDs^(62, 63)^. Interestingly, in malaria parasites the DV is an acid compartment known to be a dynamic intracellular Ca^2+^ store^(17, 64–66)^, making the subcellular location of MDR1 protein suitable for a IP_3_R-like candidate. Moreover, the *in vivo* and *in vitro* treatment with IP_3_R inhibitor 2-aminoethoxydiphenyl borinate (2-APB) is associated to reverse resistance to antimalarial chloroquine in *P. falciparum* and *P. chabaudi* parasites, presumably by disrupting Ca^2+^ homeostasis^(67)^. Multiple antimalarial drugs can also disrupt Ca^2+^ dynamic on parasite^(68, 69)^, nevertheless there is not direct evident that suggest the MDR1 acts as a Ca^2+^ gate.

Considering that agents that disrupt IP_3_R channels such as 2-APB block malaria *in vitro* growth^(9, 11, 28)^, identified this receptor in *Plasmodium* will not only add crucial missing information on malaria Ca^2+^ signalling but it will also present a potential new target for pharmacological treatment.

This work aims to stimulate the use of IP_3_-affinity column with bioinformatic strategies as a potential tool to identify proteins that might act as IP_3_R in *Apicomplexa*. The MDR1 seems to be a promising candidate to be validated, however, it is initial step from a long rewarding task of finding the *Apicomplexa* channel sensitive to IP_3_.

## Conflict of Interest

The authors declare no conflict of financial or commercial interests.

## Funding

Celia R. S. Garcia is funded by FAPESP (2017/08684-7; 2018/07177-7). Helder Nakaya is funded by FAPESP 2018/14933-2.

## Acknowledgements

We thank Prof. Dr Akio Kishigami for the helpful suggestion on the IP_3_-Affinity Column. Ross Tomaino from Taplin Mass Spectrometry Facility for helpful support on mass spectrometry data.

## Notes

### Competing Interest Statement

The authors have declared no competing interest.

