## Supplemental Table for "Combining IP_3_ affinity chromatography and bioinformatics reveals a novel protein-IP_3_ binding site on *Plasmodium falciparum* MDR1 transporter"

| Streptavidin-sepharose column | IP_3_-biotin-sepharose-streptavidin column |
| --- | --- |
| Gel section size: ~130-170kd (tracking number 32937) | Gel section size: ~130-170kd (tracking number 32941) |
| Gel section size: ~70-130kd (tracking number 32938) | Gel section size: ~70-130kd (tracking number 32942) |
| Gel section size: ~35-70kd (tracking number 32939) | Gel section size: ~35-70kd (tracking number 32943) |
| Gel section size: ~35kd (tracking number 32940) | Gel section size: ~35kd (tracking number 32944) |

**Supplement Table 1:** Table containing the tracking number of mass spectrometer database search results from *P. falciparum* protein lysate in both columns without (left) and with IP_3_ (right). The brute data is store at Taplin Mass Spectrometry, Harvard Medical School (<https://taplin.med.harvard.edu/>).

| **Gene Code** | **Predicted Function/annotation** |
| --- | --- |
| A6N5Z1_PLAFA | Helicase (Fragment) |
| ALF_PLAF7 | Fructose-bisphosphate aldolase |
| AMA1_PLAFC | Apical membrane antigen 1 |
| AN32_PLAF7 | Acidic leucine-rich nuclear phosphoprotein 32-related protein |
| ASNA_PLAF7 | ATPase ASNA1 homolog |
| C0H4C7_PLAF7 | Putative uncharacterized protein |
| C0H4M7_PLAF7 | Putative uncharacterized protein |
| C0H4X1_PLAF7 | Protein disulfide isomerase |
| C0H4Y6_PLAF7 | Protein disulfide isomerase |
| C0H529_PLAF7 | Small nuclear ribonucleoprotein (SnRNP) |
| C0H570_PLAF7 | RNA binding protein |
| C0H5G3_PLAF7 | Ribosomal protein L18 |
| C0H5I5_PLAF7 | Splicing factor 3b subunit |
| C0H5J9_PLAF7 | Putative uncharacterized protein |
| C0H5L8_PLAF7 | U2 small nuclear ribonucleoprotein A |
| C0H5U7_PLAFA | GrpE protein homolog |
| C0MHL8_PLAFA | Acid phosphatase |
| C5HX23_PLAFA | Putative erythrocyte membrane protein |
| C5HX92_PLAFA | Cysteine-rich surface protein |
| C5HXF7_PLAFA | Putative uncharacterized protein |
| C6KSP9_PLAF7 | RNA binding protein |
| C6KSR7_PLAF7 | Glutaredoxin-like protein |
| C6KST5_PLAF7 | Chaperone, putative |
| C6KSY5_PLAF7 | Putative uncharacterized protein |
| C6KT11_PLAF7 | Mitochondrial import receptor subunit tom40 |
| C6KT23_PLAF7 | 60S ribosomal protein L27a |
| C6KT75_PLAF7 | Ribonuclease |
| C6KTC6_PLAF7 | Putative uncharacterized protein |
| CDPK4_PLAF7 | Calcium-dependent protein kinase |
| HSP73_PLAFA | Heat shock 70 kDa protein PPF203 |
| IPYR_PLAF7 | Probable inorganic pyrophosphatase |
| MDR_PLAFF | Multidrug resistance protein |
| MYOA_PLAF7 | Myosin-A |
| O77309_PLAF7 | Cytoadherence linked asexual protein 3.2 |
| O77325_PLAF7 | PRP19-like protein |
| O77376_PLAF7 | Band 7-related protein |
| O77380_PLAF7 | CPSF (Cleavage and polyadenylation specific factor), |
| O77381_PLAF7 | 40S ribosomal protein S11 |
| O77395_PLAF7 | 40S ribosomal protein S15A |
| O96128_PLAF7 | Early transcribed membrane protein 2, ETRAMP2 |
| O96164_PLAF7 | Serine repeat antigen 4 (SERA-4) |
| O96191_PLAF7 | Conserved Plasmodium protein |
| O96203_PLAF7 | Peptide chain release factor subunit 1 |
| O96212_PLAF7 | Heat shock 40 kDa protein |
| O96220_PLAF7 | T-complex protein 1 |
| O96236_PLAF7 | DNA-directed RNA polymerase |
| O96259_PLAF7 | Conserved Plasmodium protein |
| O97248_PLAF7 | 40S ribosomal protein S23 |
| O97266_PLAF7 | Translation initiation factor E4 |
| PYRD_PLAF7 | Dihydroorotate dehydrogenase (quinone), mitochondrial |
| Q02602_PLAFA | ORF 2 protein |
| Q25710_PLAFA | MCP1 protein OS |
| Q689C6_PLAFA | PFC0270w protein |
| Q71KM3_PLAFA | Glyoxalase I |
| Q76NM4_PLAF7 | Rab11a, GTPase |
| Q7K6B1_PLAF7 | Protein kinase c inhibitor-like protein |
| Q7KAV0_PLAFA | Karyopherin alpha |
| Q7KQK3_PLAF7 | Heat shock protein DNAJ homologue Pfj4 |
| Q86GS5_PLAFA | Adenosine deaminase |
| Q8I0H2_PLAFA | Glutamate-rich protein (Fragment) |
| Q8I205_PLAF7 | Putative uncharacterized protein |
| Q8I207_PLAF7 | Putative uncharacterized protein |
| Q8I246_PLAF7 | Phenylalanyl-tRNA synthetase beta chain, putative |
| Q8I289_PLAF7 | Putative uncharacterized protein |
| Q8I298_PLAF7 | Putative uncharacterized protein |
| Q8I2B1_PLAF7 | Aspartyl-tRNA synthetase |
| Q8I2S6_PLAF7 | DNA-directed RNA polymerase II |
| Q8I2V9_PLAF7 | Protein disulfide isomerase |
| Q8I328_PLAF7 | Putative uncharacterized protein |
| Q8I3A2_PLAF7 | Bacterial histone-like protein |
| Q8I3C0_PLAF7 | Serine repeat antigen 9 (SERA-9) |
| Q8I3J3_PLAF7 | Ubiquitin carboxyl-terminal hydrolase |
| Q8I3M1_PLAF7 | Cytosolic preribosomal GTP-binding protein |
| Q8I3N3_PLAF7 | Mitochondrial processing peptidase alpha subunit |
| Q8I488_PLAF7 | PIESP2 erythrocyte surface protein |
| Q8I489_PLAF7 | Heat shock protein |
| Q8I492_PLAF7 | Mature parasite-infected erythrocyte surface antigen (MESA) or PfEMP2 |
| Q8I4R5_PLAF7 | Rhoptry neck protein 3 |
| Q8I4U3_PLAF7 | Conserved Plasmodium protein |
| Q8I4V8_PLAF7 | FK506-binding protein (FKBP)-type peptidyl-propyl isomerase |
| Q8I4Y5_PLAF7 | ADP-ribosylation factor GTPase-activating protein |
| Q8I546_PLAF7 | Conserved Plasmodium membrane protein |
| Q8I553_PLAF7 | Ubiquitin-activating enzyme |
| Q8I5L8_PLAF7 | Conserved Plasmodium protein |
| Q8I5P4_PLAF7 | Signal recognition particle SRP19 |
| Q8I5S6_PLAF7 | Eukaryotic translation initiation factor 3 subunit 10 |
| Q8I5Y9_PLAF7 | Histone binding protein |
| Q8I613_PLAF7 | Signal recognition particle SRP14 |
| Q8I6T2_PLAF7 | Isocitrate dehydrogenase (NADP), mitochondrial |
| Q8I6Z1_PLAF7 | Acyl-coA synthetase, PfACS5 |
| Q8IAL6_PLAF7 | Mannose-6-phosphate isomerase |
| Q8IAR6_PLAF7 | Proteasome subunit alpha type 5 |
| Q8IB14_PLAF7 | High mobility group protein |
| Q8IB51_PLAF7 | 60S ribosomal protein L22 |
| Q8IB57_PLAF7 | Small nuclear ribonucleoprotein |
| Q8IB60_PLAF7 | PfSec23 protein |
| Q8IB66_PLAF7 | RNA binding protein |
| Q8IBC0_PLAF7 | Putative uncharacterized protein |
| Q8IBC3_PLAF7 | Prohibitin |
| Q8IBE9_PLAF7 | Putative uncharacterized protein |
| Q8IBN7_PLAF7 | P36-like protein homologue |
| Q8IBQ6_PLAF7 | 60S ribosomal protein L11a |
| Q8IBR6_PLAF7 | Prefoldin subunit 3 |
| Q8IBS3_PLAF7 | Seryl-tRNA synthetase |
| Q8IBZ4_PLAF7 | Putative uncharacterized protein |
| Q8IC42_PLAF7 | Putative uncharacterized protein |
| Q8ID59_PLAF7 | DNA-directed RNA polymerase 2 |
| Q8IDB0_PLAF7 | 40S ribosomal protein S13 |
| Q8IDK3_PLAF7 | Putative uncharacterized protein |
| Q8IDP4_PLAF7 | Thioredoxin |
| Q8IDP8_PLAF7 | Aspartate carbamoyltransferase |
| Q8IDQ2_PLAF7 | Kelch protein |
| Q8IDS0_PLAF7 | Vacuolar ATP synthase subunit D |
| Q8IDV2_PLAF7 | Proteasome regulatory component |
| Q8IE02_PLAF7 | Apurinic/apyrimidinic endonuclease Apn1 |
| Q8IE10_PLAF7 | Glutaminyl-tRNA synthetase |
| Q8IE67_PLAF7 | Phosphoribosylpyrophosphate synthetase |
| Q8IE85_PLAF7 | 60S ribosomal protein L6 |
| Q8IE99_PLAF7 | Splicing factor |
| Q8IEJ5_PLAF7 | Putative uncharacterized protein |
| Q8IEK3_PLAF7 | 26S proteasome regulatory subunit 7 |
| Q8IEN3_PLAF7 | Carbamoyl phosphate synthetase |
| Q8IEQ1_PLAF7 | 26S proteasome regulatory subunit |
| Q8IHN4_PLAF7 | Antigen 332, DBL-like protein |
| Q8IHR4_PLAF7 | Dynamin-like protein |
| Q8IHU0_PLAF7 | 60S ribosomal protein L28 |
| Q8II42_PLAF7 | Conserved Plasmodium protein |
| Q8II45_PLAF7 | Ubiquitin-like protein |
| Q8II69_PLAF7 | RNA methyltransferase |
| Q8II82_PLAF7 | Conserved Plasmodium protein |
| Q8II94_PLAF7 | Small nuclear ribonucleoprotein F |
| Q8IIA4_PLAF7 | Threonine--tRNA ligase |
| Q8IIB4_PLAF7 | 60S ribosomal protein L35 |
| Q8IIC9_PLAF7 | Translation elongation factor EF-1 |
| Q8IIF6_PLAF7 | Conserved Plasmodium protein |
| Q8III3_PLAF7 | Pre-RNA processing ribonucleoprotein |
| Q8IIJ6_PLAF7 | Deubiquinating/deneddylating enzyme |
| Q8IIK5_PLAF7 | Moving junction protein |
| Q8IIQ4_PLAF7 | Actin-like protein homolog, ALP1 homolog |
| Q8IIR9_PLAF7 | Casein kinase II, alpha subunit |
| Q8IIU3_PLAF7 | RuvB DNA helicase |
| Q8IIU7_PLAF7 | Conserved Plasmodium protein |
| Q8IIU9_PLAF7 | Conserved Plasmodium protein |
| Q8IIW2_PLAF7 | Phenylalanyl-tRNA synthetase beta chain |
| Q8IJ28_PLAF7 | Antigen UB05 |
| Q8IJ37_PLAF7 | Pyruvate kinase |
| Q8IJ39_PLAF7 | Conserved Plasmodium protein |
| Q8IJ52_PLAF7 | Erythrocyte membrane protein |
| Q8IJ60_PLAF7 | Methionine-tRNA ligase |
| Q8IJ72_PLAF7 | Bromodomain protein |
| Q8IJC6_PLAF7 | 60S ribosomal protein L3 |
| Q8IJF2_PLAF7 | Conserved Plasmodium protein |
| Q8IJM0_PLAF7 | 26s proteasome subunit p55 |
| Q8IJM9_PLAF7 | Early transcribed membrane protein 10.3, etramp10.3 |
| Q8IJN9_PLAF7 | Heat shock protein 60 |
| Q8IJP3_PLAF7 | Cysteinyl-tRNA synthetase |
| Q8IJP9_PLAF7 | Transcriptional activator ADA2 |
| Q8IJW2_PLAF7 | Conserved Plasmodium protein |
| Q8IJX3_PLAF7 | RNA binding protein |
| Q8IJZ7_PLAF7 | 60S ribosomal protein L13 |
| Q8IK02_PLAF7 | 40S ribosomal protein S20e |
| Q8IK15_PLAF7 | PF70 protein |
| Q8IK70_PLAF7 | Putative uncharacterized protein |
| Q8IKA6_PLAF7 | DNAJ protein |
| Q8IKF6_PLAF7 | Putative uncharacterized protein |
| Q8IKH2_PLAF7 | Transcription factor with AP2 domain(S) |
| Q8IKH3_PLAF7 | 26S proteasome subunit |
| Q8IKQ0_PLAF7 | Putative uncharacterized protein |
| Q8IKQ5_PLAF7 | ATPase, putative |
| Q8IKT5_PLAF7 | Peptidase, putative |
| Q8IKV7_PLAF7 | Ribosome biogenesis protein tsr1 |
| Q8IL13_PLAF7 | Helicase |
| Q8IL58_PLAF7 | 60S ribosomal protein L1 |
| Q8IL71_PLAF7 | Vesicle-associated membrane protein |
| Q8ILB8_PLAF7 | Methionine aminopeptidase |
| Q8ILK1_PLAF7 | Arginine-N-methyltransferase |
| Q8ILP6_PLAF7 | Glycine-tRNA ligase |
| Q8ILR7_PLAF7 | DNA replication licensing factor MCM2 |
| Q8ILU8_PLAF7 | Ribonucleoprotein |
| Q8ILV5_PLAF7 | Putative uncharacterized protein |
| Q8IM71_PLAF7 | Choline kinase |
| Q8MTV6_PLAFA | Mitochondrial processing peptidase beta subunit |
| Q8WSM9_PLAFA | Putative uncharacterized protein |
| Q8WT07_PLAFA | Ran-binding protein |
| Q8WT10_PLAFA | ETRAMP10.1 protein (Fragment) |
| Q962H6_PLAFA | Pfs38 |
| Q962N7_PLAFA | Serine/threonine protein phosphatase PfPP5 |
| Q9GPQ6_PLAFA | Small GTPase rab11 (Fragment) |
| Q9NFE6_PLAF7 | SAC3/GNAP family-related protein |
| Q9NNW0_PLAFA | Putative uncharacterized protein (Fragment) |
| Q9U775_PLAFA | GMP synthetase |
| RBP1_PLAF7 | Reticulocyte-binding protein PFD0110w |
| RIR1_PLAF4 | Ribonucleoside-diphosphate reductase large subunit |
| RIR2_PLAF4 | Ribonucleoside-diphosphate reductase small chain |
| SAHH_PLAF7 | Adenosylhomocysteinase |
| SERA_PLAF7 | Serine-repeat antigen protein |
| SERA_PLAFD | Serine-repeat antigen protein |
| STI1L_PLAF7 | STI1-like protein |
| TBA_PLAFK | Tubulin alpha chain |
| TCPH_PLAF7 | T-complex protein 1 subunit eta |
| TOP2_PLAFK | DNA topoisomerase 2 |

**Supplement Table 2:** List of the *P. falciparum* proteins exclusively found on the IP_3_-biotin-streptavidin-sepharose column. All gene codes and their respective annotations were obtained from UniProt database ([www.uniprot.org](http://www.uniprot.org)).
